# Towards an automated protocol for wildlife density estimation using camera-traps

**DOI:** 10.1101/2024.08.05.606345

**Authors:** Andrea Zampetti, Davide Mirante, Pablo Palencia, Luca Santini

## Abstract

Camera-traps are valuable tools for estimating wildlife population density, and recently developed models enable density estimation without the need for individual recognition. Still, processing and analysis of camera-trap data are extremely time-consuming. While algorithms for automated species classification are becoming more common, they have only served as supporting tools, limiting their true potential in being implemented in ecological analyses without human supervision. Here, we assessed the capability of two camera-trap based models to provide robust density estimates when image classification is carried out by machine learning algorithms.

We simulated density estimation with Camera-Traps Distance Sampling (CT-DS) and Random Encounter Model (REM) under different scenarios of automated image classification. We then applied the two models to obtain density estimates of three focal species (roe deer *Capreolus capreolus*, red fox *Vulpes vulpes*, and Eurasian badger *Meles meles*) in a reserve in central Italy. Species detection and classification was carried out both by the user and machine learning algorithms (respectively, MegaDetector and Wildlife Insights), and all outputs were used to estimate density and ultimately compared. Simulation results suggested that the CT-DS model could provide robust density estimates even at poor algorithm performances (down to 50% of correctly classified images), while the REM model is more unpredictable and depends on multiple factors. Density estimates obtained from the MegaDetector output were highly consistent for both models with the manually labelled images. While Wildlife Insights’ performance differed greatly between species (recall: badger = 0.15; roe deer = 0.56; fox = 0.75), CT-DS estimates did not vary significantly; on the contrary, REM systematically overestimated density, with little overlap in standard errors.

We conclude that CT-DS and REM models can be robust to the loss of images when machine learning algorithms are used to identify animals, with the CT-DS being an ideal candidate for applications in a fully unsupervised framework. We propose guidelines to evaluate when and how to integrate machine learning in the analysis of camera-trap data for density estimation, further strengthening the applicability of camera traps as a cost-effective method for density estimation in (spatially and temporally) extensive multi-species monitoring programs.

## 1 INTRODUCTION

Population abundance is key parameters in ecology and conservation (Callaghan et al., 2024). The need for extensive and continuative monitoring across space and time has led to a growing employment of camera-traps as a surveying tool in the last decades (Delisle et al., 2021). Camera traps are a non-invasive, cost-effective method for monitoring mammals in a wide variety of habitats, landscapes and conditions, and near-continuously over long periods of time (Kucera & Barrett, 2011). A key advantage of camera-traps is the possibility of simultaneously collecting data over a wide range of species including nocturnal and elusive mammals, enabling multi-species surveying programs which are particularly relevant for population and community ecology studies and management efforts (Ahumada et al., 2013; O’Brien, 2010). Initially, camera-traps applications for estimating population density were limited to species that could be individually recognized. Recently, however, there has been a surge of statistical models for estimating population density without the need for individual recognition, such as Random Encounter Model (REM; Rowcliffe et al., 2008), Spatial Counts (Chandler & Royle, 2013; Evans & Rittenhouse, 2018), Camera-Trap Distance Sampling (CT-DS; Howe et al., 2017), Random Encounter and Staying Time (REST; Nakashima et al., 2018), Time-To-Event and Space-To-Event (TTE and STE; Moeller et al., 2018), Time In Front of the Camera model (TIFC; Huggard, 2018; Warbington & Boyce, 2020) and species‘ space use model (Luo et al., 2020).

Yet, applying these approaches requires intensive manual labour. Camera trapping produces substantial volume of data, which requires a considerable amount of manual work to filter relevant material and extract all key parameters for population density estimation. Consequently, much attention has been directed towards the development of machine learning algorithms (specifically, *deep learning*; Wäldchen & Mäder, 2018) for automated image processing (Vélez et al., 2023). These algorithms are trained on vast pre-processed datasets to perform specific tasks that would otherwise be performed manually (Green et al., 2020). Examples include the removal of images not containing animals (Beery et al., 2019), counting the number of individuals (Norouzzadeh et al., 2018), and species (Rigoudy et al., 2023; Tabak et al., 2020) or individual animal identification (Cheema et al., 2017). Despite the variety of machine learning models developed to support data processing in camera trapping studies, their implementation in real monitoring contexts remains largely unexplored. Most published studies have evaluated their performance in different ecological contexts to assess their transferability (Tabak et al., 2020; Vélez et al., 2023). Several studies showed suboptimal precision parameters in out-of-sample images or videos (Vélez et al., 2023), concluding that a fully automated approach is currently beyond the capabilities of most algorithms. A minority of studies have compared estimates of ecological parameters obtained from data processed by operators and machine learning algorithms (Gimenez et al., 2021; Mitterwallner et al., 2023; Whytock et al., 2021), but the bias resulting from suboptimal performance of these algorithms in density estimation models has not yet been quantified.

Many of these models are based on the estimation of detectability and the encounter rate between animals and camera-traps (Palencia, Rowcliffe, et al., 2021). Assessing how the imperfect detection of machine learning algorithms (i.e., animals captured by cameras but not identified by the algorithm) interacts with the intrinsic imperfect detection of camera traps (i.e., animals present in cameras‘ field of view but not captured) can provide important insights on the implications of an automated image classification approach for animal population density estimation. A number of factors can influence how automated image classification errors can propagate into the final density estimates. For example, missing animals due to misclassification can result in a lower capture rate and in an underestimation of density. On the other hand, if faster moving animals are more likely missed due to motion blur, movement speed can be underestimated hence inflating density estimation. Thus, an assessment of how these parameters concurrently change and interact under an automated image classification scenario is needed.

In this paper, we test the robustness of a semi-automated approach for density estimation through camera-traps. We focus on CT-DS and REM models as they are the most commonly used and best described frameworks for density estimation with camera-traps (Gilbert et al., 2021). First, we present a simulation to explore the interaction between different parameters and sources of errors on CT-DS and REM performance. Then, we present a real case study on three focal species in central Italy (roe deer *Capreolus capreolus*, Eurasian badger *Meles meles*, red fox *Vulpes vulpes*). We evaluate machine learning performance in an automated species detection and classification framework, and assess the effect of data loss due to suboptimal algorithms performance on the reliability of final density estimates.

## 2 MATERIALS AND METHODS

### 2.1 Simulations

A number of factors can influence how the error of automated image classification can propagate into the final density estimates causing underestimation or overestimation of the parameter, thus hampering our understanding of the underlying mechanism. To unveil actual operating mechanisms, we simulated the whole process by varying parameters to assess their effect on the results. We set 20 virtual camera-traps to be active for 30 days. We then simulated the detection process directly at camera level. Camera-trap detection ability was varied to account for different camera models and survey conditions: the distance of maximum capture probability was varied between 1 m and 5 m, and the decay of the detection function was modelled after a half-normal curve with 2 < σ < 5 (where σ represents the scale parameter that determines the rate at which detection probability decreases with distance). The extent of the camera‘s field of view (FOV) was set at 0.9599 radians (55°). We constrained the maximum distance at which animals could be captured to 15 m. We varied animal activity rate (i.e., the proportion of the day in which animals are detectable by cameras; Howe et al., 2017; Nakashima et al., 2018) between 0.20 and 0.60, and animal movement speed was extracted from a log-normal distribution (−0.3 < μ < 0, σ = 0.3; μ and σ represent the mean and standard deviation of the natural logarithm of the speed, respectively) to account for a broad range of target species. We did not consider animals moving in groups. For each successful capture event, we retained animal distance and angle from the camera and its movement speed. Then, we simulated the imperfect detection of a machine learning algorithm for species classification on the dataset produced. We considered four plausible shapes for the algorithm‘s detection function (uniform, linear, half-normal and hazard-rate) to simulate the missed classifications of the machine learning classifier at further distances (Figure S4), and for each we varied the intercept (which represents the probability of detection at zero distance from cameras) between 0.5 and 1, and the decay with distance to result in between 0.9 and 0.1 capture probability at 15 m. Also, we penalized detection probability for faster-moving animals according to a linear function with slope between -0.0067 and -0.0600, to simulate the loss of recognized animals due to motion blur (as suggested by the field study; see Figure S5). The simulation produced two dataset, one reflecting camera-traps performance, and one camera-traps and machine learning algorithm performance, which will be later used for the two estimation methods described in sections 2.1.1 and 2.1.2. Each 30-days survey was replicated for 10.000 iterations.

#### 2.1.1 Camera-Trap Distance Sampling (CT-DS)

CT-DS is the adaptation of the distance sampling workflow to camera-traps, considering cameras as observers on point transects (Howe et al., 2017). Population density estimation (D) is estimated as:

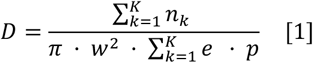

where *e* is the sampling effort expended at the point *k* calculated as 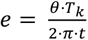, *K* is the set of points, *θ* is the total width of the FOV of camera-traps, *T*_*k*_ is the sampling period at point *k, w* is the truncation distance, *n*_*k*_ is the number of animal observation at the point *k*, and *p* is the probability of capturing an animal within θ and *w*. As cameras sample continuously in time, the *t* parameter is used to divide total sampling effort in discrete intervals. For the simulations, we considered t = 1 s and we varied the average encounter rate between 0 and 20 captures/day. We fitted distance sampling models on the retained animal distances of both the original and machine learning-filtered datasets to estimate the effective detection radius using the ‘Distance‘ package (Miller & Clark-Wolf, 2022) in R (R Core Team, 2023). For the detection function, we considered all the combinations between the half-normal, hazard-rate, and uniform families, using up to the first adjustment term from each series: cosine, simple polynomial, and Hermite polynomial. As longer distances carry little information and may cause the detection function to be heavy-tailed (Buckland et al., 2001), we right-truncated distances at the 95^th^ percentile when detection probability was lower than 0.1 (Howe et al., 2017; Palencia, Rowcliffe et al., 2021). Model selection was conducted following the QAIC procedure indicated by Howe et al. (2019) for overdispersed data, using functions from the ‘Distance‘ package (Miller & Clark-Wolf, 2022). Finally, density was compiled through the ‘Distance‘ package following equation [1] and multiplying the estimate for the inverse value of the proportion of the area sampled by cameras (i.e., θ/2*π*) and the inverse value of the activity level. For each iteration, we produced two separate density estimates: one from the original dataset and one from the dataset where we simulated the automated classification.

#### 2.1.2 Random Encounter Model (REM)

REM is based on the modelling of animals capture rate by the cameras by taking into account animal movement parameters and cameras‘ detection abilities (Rowcliffe et al., 2008). Population density (*D*) is calculated as:

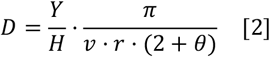

where *Y* is the number of independent animal capture events, *H* is the total survey effort across all camera stations, *v* is the average daily distance travelled by the animal (hereafter referred to as “day range”), while *r* and θ describe the radius and angle of camera-traps effective detection zone. We repeated the simulations as described for CT-DS, but this time for the machine learning-filtered dataset we also modelled random animal movement in front of the cameras: for simplicity, we did not consider path tortuosity and assumed that the animals would cross the FOV following a straight line. We varied the average encounter rate between 0 and 5 independent sequences/day, then we simulated animal passes as a series of snapshots to which we applied the machine learning filter. For each encounter, speed of movement, distance and angle from the first snapshot successfully captured (i.e., first image in each capture sequence) were retained. We identified animal speed classes based on different movement behaviours using the ‘trappingmotion‘ package (Palencia, 2023) in R and we estimated day range as the sum of the product of the mean speed and the proportion of activity associated with each behaviour class (Palencia, Fernández-López, et al., 2021). Methods, R packages, and practical considerations about detection functions and truncation decisions were the same described previously for the CT-DS model (see 2.1.1). We finally estimated density using functions from the ‘camtools‘ package (Rowcliffe, 2019) in R. Overall variance was estimated using the delta method (Palencia, Rowcliffe, et al., 2021; Rowcliffe et al., 2008; Seber, 1982), in order to consider the uncertainty associated with each parameter (*v, r* and *θ*; equation [2]). For each iteration, we produced two separate density estimates from the original dataset and the one where we applied the automated classification.

#### 2.1.3 Analysis of simulation output

To unveil the contribution of the different parameters and their interaction to the estimated density from the two datasets, we applied a random forest algorithm to the simulated data using the ‘randomforest‘ package (Cutler & Wiener, 2022). We used the relative difference in density between the user and AI estimates as response variable, calculated as (density_AI_ - density_user_) / density_user_, and the following AI parameters as predictor variables: the shape of the AI algorithm‘s detection function (*AI_det*); the intercept of the AI algorithm‘s detection function (*intercept*), which defines the detection probability at distance = 0 m; the heaviness of the tail of the AI algorithm‘s detection function (*prob_max*), which defines the detection probability at distance = 15 m; the *recall* metric; and only for REM, the slope of the penalizing factor for animal speeds (*speed_det_slope*). For both models, we built a forest of 1000 trees, and we tuned the model by setting the number of variables sampled at each split (*mtry* parameter) from 2 to 4 (Hastie et al., 2009), and selected the one resulting in the lowest error. We assessed variable importance and marginal effects using functions from the ‘pdp‘ (Greenwell, 2022) package in R.

### 2.2 Case study

We conducted the data collection in the Tenuta Sant‘Egidio, a private reserve of approximately 130 ha located on Mount Cimino near Viterbo in central Italy (42.417 N, 12.219 E; Figure S1). The area is covered by Mediterranean wood dominated by chestnut (*Castanea sativa*) and oak trees (*Quercus* spp.), and ranges between 450 and 750 m altitude. We employed a total of 20 Browning Patriot camera-traps from August to September 2022. Cameras were placed along the intersection of a systematic grid with random origin and 250 m spacing. We considered a 10 m buffer around the generated positions in order to mount cameras on suitable trees and avoid very unfavourable conditions for animal detections (e.g., the middle of dense shrubs). For two cameras the sampling points were inaccessible, so they were placed in the nearest available locations (approximately 100 m from the original spot). All camera-traps were placed at 60-80 cm from the ground, with the FOV starting at 1.50 m from the camera, angled to be parallel to the slope of the ground, and facing North to avoid light over-exposition. Cameras were not baited. To ensure spatial movement resolution for fast-moving animals, we set cameras to record three rapid fire burst images (0.15 s delay), and with the minimum delay period possible between bursts (1 s). During the installation of each camera, we placed markers in the FOV with 0.50-1.50 m spacing (depending on the slope of the ground) that were later imported in the software Adobe Photoshop 2023 v.24.0.0 to layout a virtual grid. This was later used to estimate animal distances from cameras and movement parameters needed for the two population density estimation methods.

### 2.3 Species classification

First, we cleaned up the data with the aid of a machine learning algorithm for animal detection and produced two datasets, one accepted as it is and one corrected for false positives (2.3.1). Subsequently, we labelled photos of the corrected dataset at the species level manually, and those of the uncorrected dataset using another machine learning algorithm for species classification, thus producing a third dataset (2.3.2). Finally, we estimated classification performance of the two algorithms (2.3.3).

#### 2.3.1 Manual classification with MegaDetector

We carried out the first images classification process in a semi-automated (“*man-in-the-loop*”) workflow, in which the output of a machine learning algorithm for animal detection was used solely as a visual aid for the user to speed up the process (Vélez et al., 2023). We used the MegaDetector algorithm (Beery et al., 2019), hosted at the agentmorris/MegaDetector GitHub repository. The model is trained on an extensive global dataset and can pre-process camera-trap images in broad categories (“animal”, “person”, “empty” and “vehicles”) to facilitate the sorting process by the user (Greenberg, 2020).

For this study, we used the MegaDetector v5a model through the EcoAssist platform (Van Lunteren, 2023). We implemented a post-processing step provided in the MegaDetector GitHub repository (see agentmorris/ MegaDetector/megadetector/postprocessing/repeat_detec tion_elimination) through a Python script to reduce the number of false positives (e.g., logs, rocks or branches resembling parts of animals). We then analysed the output in the Timelapse2 v.2.3.0.0 software (Greenberg & Godin, 2012), which enables the display of bounding boxes around the detected animals for visual aid and lets the user filter for different confidence levels from the MegaDetector classification. In this study, we used the default threshold settings. Using this pipeline, we were able to first analyse the images labelled as “empty/person/vehicle” by the algorithm, correct for eventual false positives, and quickly discard all these images from the dataset. Then, the images classified as “animals” were inspected with the help of the bounding boxes drawn by MegaDetector and classified at the species level directly on the Timelapse2 interface thanks to the built-in metadata labelling option. A separate dataset (hereafter, MD-dataset), not corrected for false negatives (i.e., images of a given species missed or misclassified by the algorithm), was retained to later be used directly in the analyses.

#### 2.3.2 Unsupervised classification with Wildlife Insights

To evaluate the effects of a fully automated image classification on density estimates, we selected a second machine learning algorithm for species identification. We chose Wildlife Insights, a web-based platform (https://www.wildlifeinsights.org) developed by an international partnership (Ahumada et al., 2020). The initiative serves as a data library and data-sharing platform in the cloud (Vélez et al., 2023), where users can upload their camera-trap images for species classification, preview and correct/annotate the labels on the cloud and download results and image metadata. The choice of the platform was motivated by the extensive training dataset for the algorithm (>35 million images for 1295 species and 237 higher taxonomic classes; information updated as of August 2024). The platform is open access and does not require informatic or coding skills to conduct automated species classification. Furthermore, classification results can be filtered based on different hierarchical taxonomic levels (i.e., species, genus, family, order, class), thus enabling versatility in interpreting results.

For this study, camera-trap images were uploaded to the Wildlife Insights platform for taxonomic identification. Classification results were then downloaded and treated as an independent dataset with respect to the user-classified images. We decided to consider classification at the family level for the roe deer (as no other species from the Cervidae family occur in the study area). After an exploratory analysis, the same decision was made for the red fox due to the extremely low number of wolf images (n = 25, distributed over 2 capture events) that would have been included. The higher taxonomic levels were used to ensure the highest predictive ability possible from the algorithm. Finally, badger classifications were considered at the species level. The Wildlife Insights output (hereafter, WI-dataset) was not corrected for false negatives by the user and was used directly in the analyses.

#### 2.3.3 Evaluation of algorithms performance

We evaluated the performance of MegaDetector for animal detection and Wildlife Insights for animal classification by comparing the algorithms‘ output with the user manual classification. We used functions from the ‘caret‘ package (Kuhn, 2019) in R to build a confusion matrix for the observed and predicted classes, and then calculate model precision [3] and recall [4] as metrics of true positives and false negatives rates (Sokolova & Lapalme, 2009). Precision [3] is calculated as:

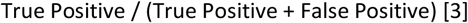

and expresses the proportion of the correctly predicted classifications over the total predictions. Recall [4] is calculated as:

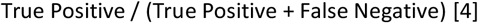

and expresses the proportion of the correctly predicted classifications over the actual number of animals present.

### 2.4 Density estimation

We then estimated population density using CT-DS and REM models on the three datasets: the user-dataset, the MD-dataset and the WI-dataset.

#### 2.4.1 Camera-Trap Distance Sampling (CT-DS)

To quantify the portion of the day in which animals were active, we estimated activity level of each target species following Rowcliffe et al., (2014) using the ‘activity‘ package (Rowcliffe, 2019) in R. We extracted animal distances from cameras by overlaying the virtual grid to images, considering all snapshots corresponding to *t* = 1 s intervals in order to maximize sample size. We then followed the same procedure described for the simulations (see 2.1.1) to estimate density, but this time we considered up to the second adjustment terms for all the combinations of the distance sampling detection function. We also left-truncated data at the 5^th^ percentile to account for animal passing beneath the cameras at shorter distances (Buckland et al., 2001). Moreover, right-truncation for roe deer in the user-classified dataset caused failure in computing confidence intervals, so we decided not to right-truncate distances in that case. While methods for avoiding bias due to animal reactions to camera (e.g., curiosity or alarm) have been described (Delisle et al., 2023), our dataset included very few detections where individuals displayed attractive behaviour (less than 3% of the total encounters), so we did not expect them to cause bias. As the assumption of independency of captures is violated due to the consideration of multiple detections of the same animal, we estimated variance using a non-parametric bootstrap with replacement (n = 1000) between camera-trapping sites (Buckland, 1984; Buckland et al., 2001; Howe et al., 2017). The same process was replicated for both the user-dataset, and the MD- and WI-dataset to obtain separate parameters and density estimates.

#### 2.4.2 Random Encounter Model (REM)

Following Rowcliffe et al. (2008), we considered an individual entering and exiting the FOV as an independent encounter. Animal activity level was estimated with the same procedure described for CT-DS. We calculated average animal speed by overlaying the virtual grid to images, measuring distance travelled in each encounter and dividing by the duration of the sequence. For speed measurements, we discarded all the encounters where animals showed reactions to cameras, and we removed a single encounter of roe deer where the animal rested in front of the camera for more than 40 minutes. We then followed the same procedure described for the simulations (see 2.1.2) to estimate density, and as in CT-DS we left-truncated data at the 5^th^ percentile, and we considered up to the second adjustment terms for all the combinations of the distance sampling detection function. The same process was replicated for both the user-dataset, and the MD- and WI-dataset to obtain separate parameters and density estimates.

## 3 RESULTS

### 3.1 Simulations

For CT-DS, the best random forest model was obtained by sampling 2 variables at each split, and had a variance explained = 77.48%, and a Mean Squared Error (MSE) = 0.006. The most important predictor was the algorithm‘s ability to classify animals at zero distance (*intercept*), followed by *recall* and the heaviness of the tail of the AI detection function (*prob_max*) (Figure S2a). Partial dependence plots showed a distinct negative relationship between the relative variation in density and both *intercept* and *recall*, and no relationship with *prob_max* (Figure 1). The interaction between *intercept* and *recall* resulted in a compensating effect on the variation in density, where the underestimation of the parameter was minimal for high values of the two predictors (Figure 3a). For REM, we achieved the best random forest model performance with 4 variables sampled at each split, explaining 16.47% of variance and with a MSE = 0.013. The most important predictor was the *recall*, followed by the algorithm‘s ability to classify animals at zero distance (*intercept*) and the heaviness of the tail of the AI detection function (*prob_max*) (Figure S2b). The variation in density showed to be constant down to *recall* values of around 0.15, where it showed a sudden drop. Similar to CT-DS, *intercept* exhibited a negative relationship with the variation in density (although less pronounced), and no relationship with *prob_max* (Figure 2). Contrarily to CT-DS, no interaction between *intercept* and *recall* was observed, and the variation in density displayed a more independent relationship with the two predictors (Figure 3b).

**FIGURE 1.**
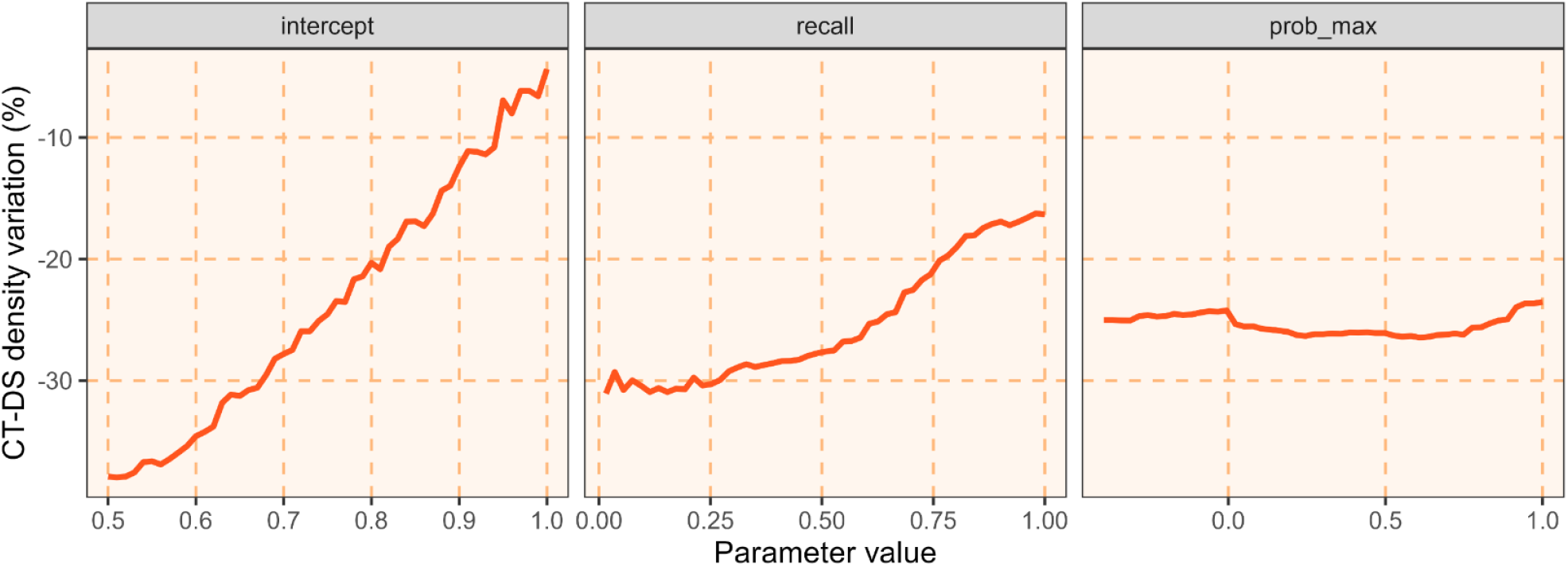
Marginal effects on the relative variation in density for the three most important predictors from the random forest model fitted on the CT-DS simulation: a) *intercept* (AI detection probability at distance = 0), b) *recall* (proportion of the correctly predicted classifications over the total number of detections), c) *prob_max* (heaviness of the tail of the AI detection function).

**FIGURE 2.**
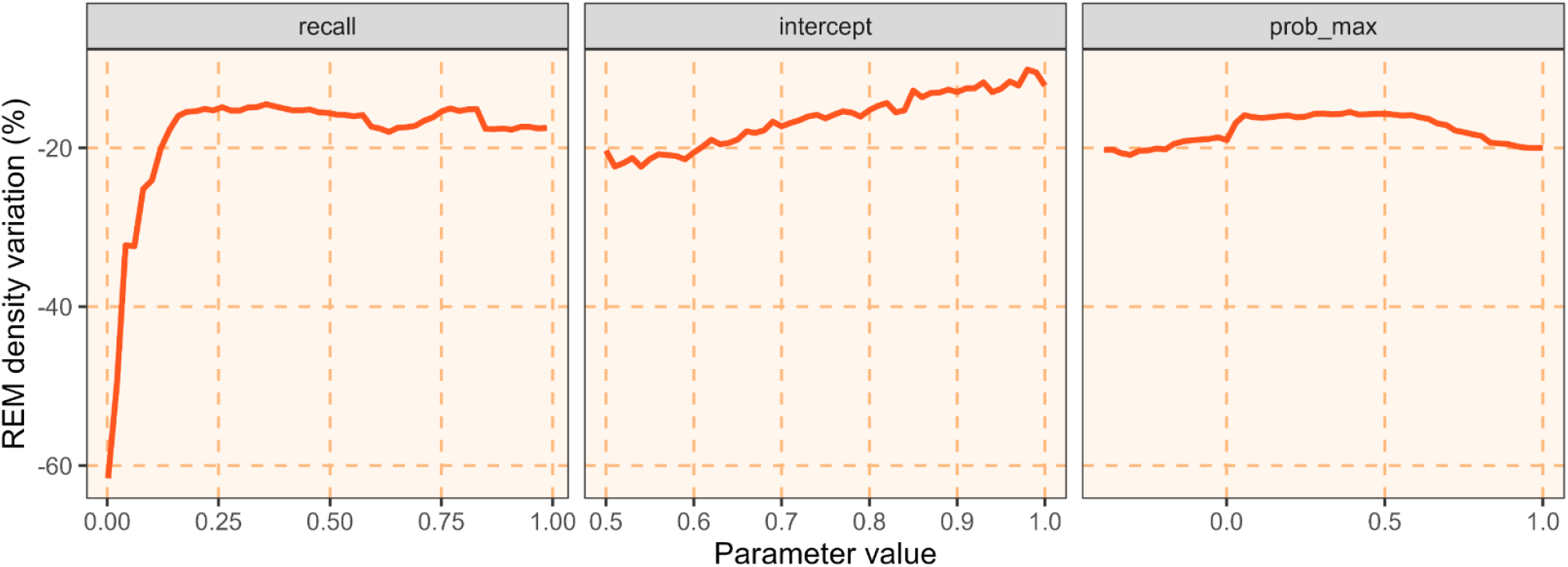
Marginal effects on the relative variation in density for the three most important predictors from the random forest model fitted on the REM simulation: a) *recall* (proportion of the correctly predicted classifications over the total number of detections), b) *intercept* (AI detection probability at distance = 0), c) *prob_max* (heaviness of the tail of the AI detection function).

**FIGURE 3.**
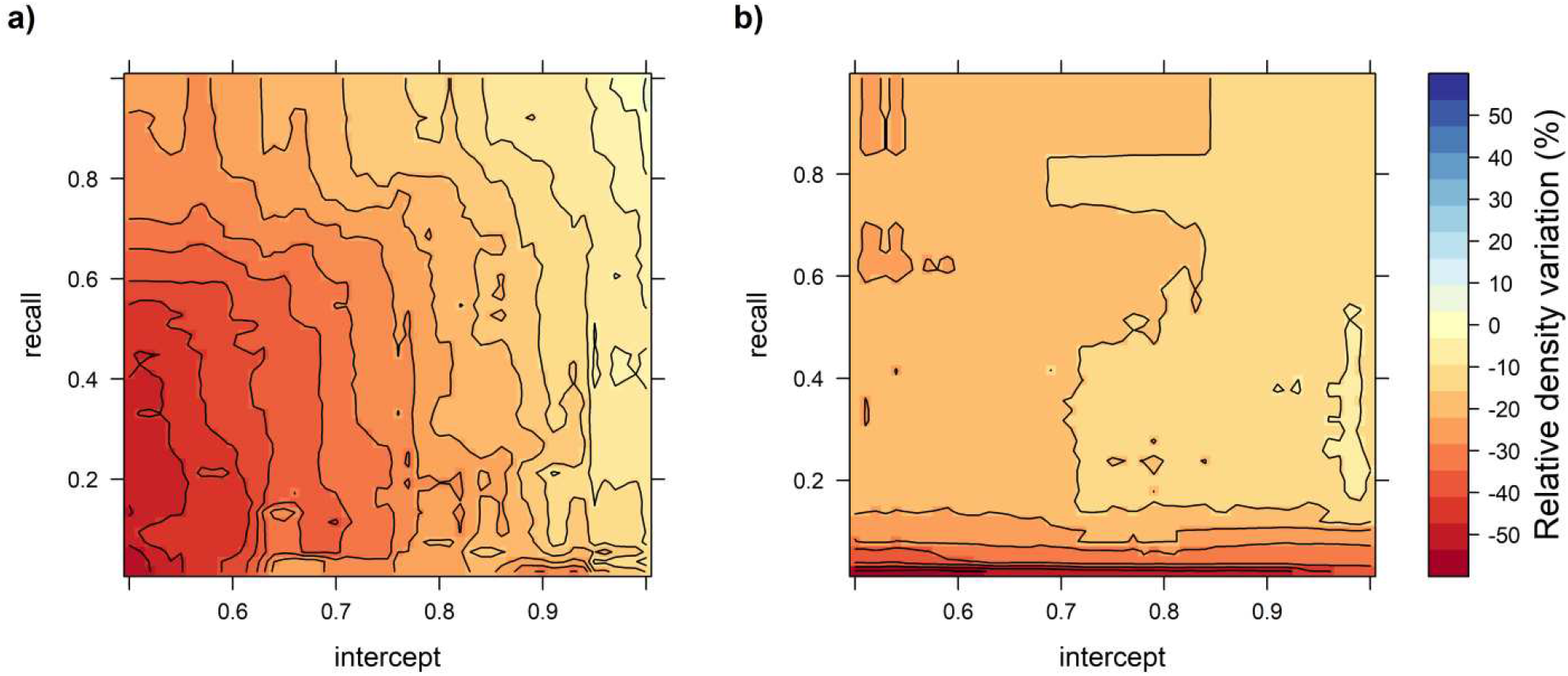
Two-ways partial dependence plots on the variation in density of the two most important predictors in the random forest models for a) CT-DS and b) REM. Regions nearer to red on the colour scale indicate underestimation of the density parameter, while blue indicates overestimation. Yellow regions show compensation between parameters that results in less biased estimates, thus better performances of the model in estimating unbiased density after the drop in images due to AI.

### 3.2 Case study

The field survey resulted in a total of 30,217 images from 19 camera-traps (due to one camera being stolen) over a period of 39 days (from August 8, 2022, to September 15, 2022; cumulative survey effort: 727 sampling days). These images were categorized into 18,626 images with animals, 6,955 images with people, and 4,636 empty images. For the target species of this study, 776 images of badger, 1,053 images of fox, and 1,293 images of roe deer were identified.

### 3.3 Species classification

MegaDetector was comparatively more prone to omission (missed detections) than commission (erroneous detections) error on animal images (precision = 0.98, recall = 0.90), and after the post-processing a further number of false positives (113 images) was successfully removed. Since the implementation of CT-DS and REM requires manual visualization of target snapshots for the extraction of distances, angles, and movement speed parameters, therefore allowing for identification and elimination of false positives, we will now focus on the recall parameter as a metric of false negative rate. Species classification results by Wildlife Insights varied greatly among focal species: the best predictions were for roe deer (recall = 0.75), followed by red fox (recall = 0.56), with Eurasian badger being the worst recognized animal by the algorithm (recall = 0.15) (Figure S5).

### 3.4 Density estimation

#### 3.4.1 Camera-Trap Distance Sampling (CT-DS)

Density estimations obtained from the user-classified dataset resulted in 1.31 ± 0.87 ind/km^2^ for the Eurasian badger, 1.94 ± 0.83 ind/km^2^ for the red fox, and 1.09 ± 0.57 ind/km^2^ for the European roe deer. Comparatively, densities estimated from the machine learning outputs resulted in 1.30 ± 0.86 ind/km^2^ for the Eurasian badger, 2.08 ± 0.80 ind/km^2^ for the red fox, and 1.13 ± 0.57 ind/km^2^ for the European roe deer (MD-dataset), and 0.27 ± 0.17 ind/km^2^ for the Eurasian badger, 2.64 ± 0.85 ind/km^2^ for the red fox, and 1.20 ± 0.64 ind/km^2^ for the European roe deer (WI-dataset) (Figure 4). A summary of all model parameters obtained for the analyses can be found in Table S1.

**FIGURE 4.**
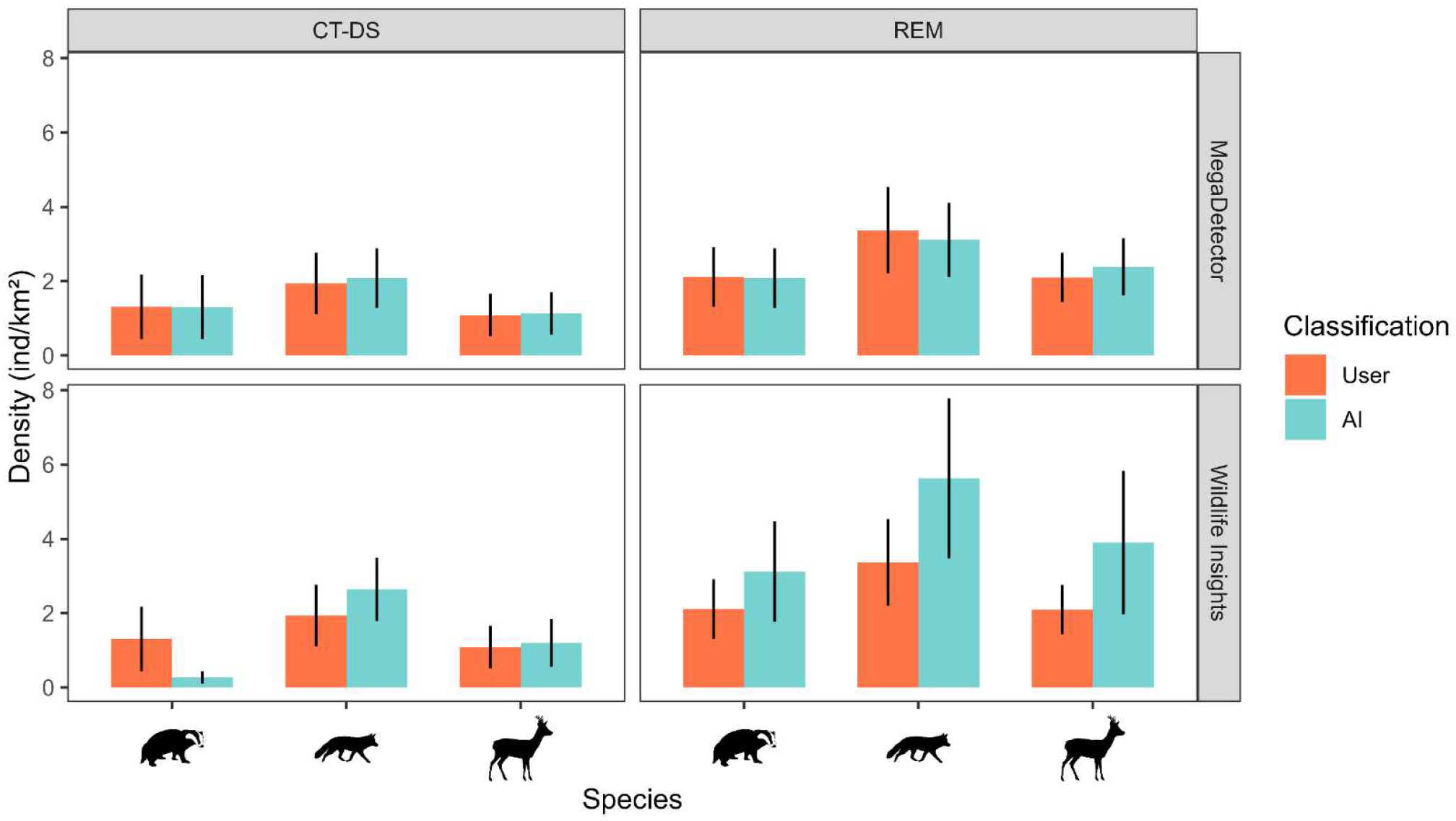
Density estimations for *Meles meles, Vulpes vulpes* and *Capreolus capreolus*, plotted in pairs between the user- and AI-datasets (MegaDetector and Wildlife Insights), both for Camera-Trap Distance Sampling (CT-DS) and Random Encounter Model (REM). For each estimate, the relative Standard Error is shown.

#### 3.4.2 Random Encounter Model (REM)

Density estimations obtained from the user-classified dataset resulted in 2.11 ± 0.80 ind/km^2^ for the Eurasian badger, 3.37 ± 1.16 ind/km^2^ for the red fox, and 2.10 ± 0.66 ind/km^2^ for the European roe deer. Comparatively, densities estimated from the machine learning outputs resulted in 2.08 ± 0.80 ind/km^2^ for the Eurasian badger, 3.11 ± 1.00 ind/km^2^ for the red fox, and 2.38 ± 0.77 ind/km^2^ for the European roe deer (MD-dataset), and 3.12 ± 1.35 ind/km^2^ for the Eurasian badger, 5.63 ± 2.15 ind/km^2^ for the red fox, and 3.90 ± 1.93 ind/km^2^ for the European roe deer (WI-dataset) (Figure 4). A summary of all model parameters obtained for the analyses can be found in Table S1.

## 4 DISCUSSION

Manual processing of data generated by camera traps can hinder their applicability density estimates for wide-scale monitoring (Ahumada et al., 2020). Machine learning models for species detection and classification have the potential to alleviate this burden, but the effect of error propagation on the final estimates has not yet been explored. Our results show that both CT-DS and REM are relatively robust to missed detections, while CT-DS is also relatively robust to misclassifications.

Although it is reasonable to expect that the loss of images would have a clear impact on density (equations [1] and [2]), simulation results showed that the recall parameter (measuring robustness to omission errors) does not always yield a strong effect on the density estimates. In fact, the interaction between recall and other significant parameters can result in compensation effects that mitigate the bias. With CT-DS, the most important predictor of density estimates was the algorithm‘s ability to classify animals at zero distance, then followed by the recall. When the algorithm correctly classified all animals close to the camera (intercept = ∼1) and the recall was higher than 0.5, the relative change in the estimated density was contained between 0 and -10% (Figure 3a). This is coherent with the distance sampling assumptions that all animals on the transect line are detected, and deviations from this assumption result in biased estimates (Buckland et al., 2015). Even if the algorithm‘s performance is suboptimal in our case, the compensating effects take place as long as detection probability at 0 distance approaches 1. Considering that the algorithm‘s classification ability decreases with distance, the detection probability at zero distance and the recall values are closely bounded: to achieve relatively high values of recall the intercept needs to be close to 1, since most missed detections will be at higher distances. Thus, when evaluating algorithms‘ performance in a density estimation scenario, values of recall down to a certain threshold would imply model‘s resilience to the loss of images, resulting in reliable density estimates. We emphasize that in a real-world scenario, this behaviour can be detected early in the exploratory analyses of the algorithm‘s performance. By assessing the recall metric on a subsample of the full data, it is possible to estimate the classification performance of individual species, thus anticipating how the model will react on a case-by-case scenario. The shape of the algorithm‘s detection function also had some minor influence on the reduction in density (Figure S3), with the half-normal and hazard-rate curves resulting in a slightly higher compensating effect. This is coherent with our expectations, since in those cases the algorithm would behave exactly as expected from a human observer with imperfect detection, where the drop in detections would be accounted for by the detection function (Buckland et al., 2001). In contrast, in REM, irrespective of the recall value, there was a consistent underestimation of the density parameter. The REM model‘s consistent response to image loss is reasonable because it relies on sequences of animal detections rather than individual capture events. This approach inherently makes the model more robust to suboptimal detection performance (Palencia, Rowcliffe, et al., 2021). However, no compensating effect due to parameters interaction emerged, and a general tendency of the model to underestimate density was observed. This contrasted with our findings in the field study, which showed the opposite trend for all three focal species (although standard errors overlap; Figure 4). So, while it is possible to advance reasonable hypotheses on the CT-DS model behaviour based on the performance of the machine learning algorithm alone, it is still unclear how image classification through machine learning influences the REM model outcome. Given the relevance of population density and abundance in the assessment of threatened species (IUCN Red List, criteria A, C, and D; IUCN, 2024) and in the application of hunting and culling quotas (Gortázar & Fernandez-de-Simon, 2022), we stress that it is particularly risky to rely on methods that could mistakenly yield higher estimates. We conclude that the REM model is robust to missed detections but not to misclassifications, hence it is currently unsuitable for fully automated approaches where both animal detection and taxonomic classification are carried out by machine learning algorithms.

Interestingly, we found CT-DS to produce comparatively lower estimates than REM, in all classification scenarios, a tendency already documented in previous studies (Corlatti et al., 2020; Palencia, Rowcliffe, et al., 2021). This might originate from sub-optimal performance of camera-traps, where the delay time between burst of images and the number of actual images in a burst deviate from the declared manufacturer settings, thus resulting in higher missed detections leading to density underestimation (Palencia, Rowcliffe, et al. 2021). For the purpose of this work, it is interesting to note that the differences between models were far greater than the differences between manually- and automatically-classified images, suggesting that the deviation from “true” density derived from an automated approach is contained enough to make reliable density estimates. This was true both in CT-DS and REM when MegaDe distances, angles and movement parameters of captured animals. At present, two approaches appear promising avenues to fully automate density estimation. The first one leverages monocular depth estimation algorithms to estimate distances based on image calibration: this has already been proposed for camera-traps (Haucke et al., 2022), and it was shown that it can be a feasible implementation in CT-DS model with minimal bias compared to manually obtained distances (Henrich et al., 2024). In conjunction with our approach and with little adjustment in survey design, this could already make the CT-DS model an ideal candidate for fully automated density estimations. The REM model poses more challenges since, in addition to distances and angles, animal paths are also required to compute day range estimates: a more sophisticated tool would be needed to project and track animal movements with respect to the ground plane in front of the camera. The second approach consists in deriving animal distances through a photogrammetric method. While this has already been described (Cui et al., 2020; Leorna et al., 2022) with successful integrations in CT-DS (Palencia et al., 2024; Zuleger et al., 2022) and REM (Palencia et al., 2023), it still implies considerable human effort to pre-process the images, and, in some cases, knowledge about the focal species‘ body traits (e.g., height at the withers or body length) that need either to be obtained from field studies or extracted from the literature (Cui et al., 2020). Future research avenues might explore the photogrammetry approach in relation to the outputs of object detection algorithms (e.g., MegaDetector): since those produce bounding boxes around the detected animals, identifying a way to reliably relate the height/width/area of the bounding boxes to animal distance from cameras could enable the extraction of the measures of interest directly from the output of the machine learning algorithm used to recognize animals.

This study stems from the necessity of developing new and efficient protocols for gathering population density data over large scales to address the “Prestonian shortfall” (i.e. lack of knowledge about the abundance of species in space and time; Hortal et al., 2015). Such a goal is strongly limited both by the costs related to obtaining accurate estimates, and by the rapid fluctuations in population abundances that require frequent and repeated assessments (Hortal et al., 2015). Implementing large-scale standardized survey protocols for long-term monitoring requires a sustainable trade-off between costs and benefits. Initiatives such as TEAM (Meek et al., 2014), Wildlife Insights (Ahumada et al., 2020), Snapshot USA (Shamon et al., 2024), and European Observatory of Wildlife (EOW; https://wildlifeobservatory.org) are fitting examples of efforts aimed at extending camera-trapping protocols beyond the sole purpose of local management. Wildlife Insights and EOW, in particular, represent cloud-based platforms with a strong emphasis on storing and sharing camera-trap data across the globe. While both initiatives already started integrating machine learning tools for automated species classification, ecologists are still reluctant to trust these tools enough to use their output directly in ecological analyses without human verification. Here we show that machine learning algorithms for animal detection can be safely integrated into density estimation workflows, and under certain circumstances, species classification can be used as a further step towards automatization. We conclude that machine learning should be increasingly considered as a key supporting tool for camera-trapping networks, enabling rapid and reliable estimates of animal density and abundance.

## AUTHOR CONTRIBUTIONS

Andrea Zampetti, Davide Mirante and Luca Santini conceived the ideas and designed the methodology in consultation with Pablo Palencia. Andrea Zampetti and Luca Santini designed the simulations; Andrea Zampetti and Davide Mirante collected the field data. Andrea Zampetti analysed the data and led the writing of the manuscript. All authors contributed critically to the drafts and gave final approval for publication.

## AKNOWLEDGEMENTS

Pablo Palencia received support from the University of Oviedo through a Juan de la Cierva contract JDC2022-048567-I supported by “Ministerio de Ciencia e Innovación”, “Agencia Estatal de Investigacion” and “NextGeneration EU” (MCIN/AEI/10.13039/ 501100011033). All the authors thank the staff of Tenuta Sant‘Egidio for their precious field support and collaboration for data collection.

## DATA AVAILABILITY STATEMENT

Data and code available from the Zenodo repository at https://doi.org/10.5281/zenodo.13152411 (Zampetti, 2024).

## SUPPORTING INFORMATION

Additional supporting information can be found online in the Supplementary Material section.

**Figure S1.** Tenuta Sant‘Egidio reserve in Mount Cimino, Viterbo, central Italy. The black dots represent the locations of camera-traps within the study area.

**Figure S2.** Predictive importance of each variable in the random forest models according to the percentage increase in Mean Squared Error (MSE) for a) CT-DS and b) REM. Higher values of %MSE increase are associated with a higher predictive importance of that particular variable in explaining the variation in density when a machine learning algorithm is used to classify images.

**Figure S3.** Marginal effects on the relative variation in density for the different shapes of the AI detection function considered in the simulations. The half-normal and hazard-rate curves exhibit a slightly better compensation effect, resulting in less underestimation of the density parameter.

**Figure S4**. Proportion of observations correctly captured by Wildlife Insights by distance for each focal species. For each class, the proportion is calculated as the number of captured observations over the total number of observations in that class.

**Figure S5.** Proportion of observations missed by Wildlife Insights by speed class for each focal species. For each class, the proportion is calculated as the number of missed observations over the total number of observations in that class.

**Figure S6.** Confusion matrices for model performance of a) MegaDetector (animal detection) and b) Wildlife Insights (animal classification). Cells show the proportion of correct classifications for each class against the ground truth, allowing visualization of exclusion errors for that class. “Other” refers to the non-focal animal species captured during the camera-trapping survey.

**Table S1.** Estimated models‘ parameters and relative standard errors for REM and CT-DS for the three focal species, both using the user-classified dataset and the WI (Wildlife Insights)-classified dataset. Note that for CT-DS the total capture events differ from the total number of images for that species, due to the fact that t = 1 s was considered for the snapshots so if multiple images were recorded within the same second, only the first was selected.

