## supplementary_material for "Towards an automated protocol for wildlife density estimation using camera-traps"

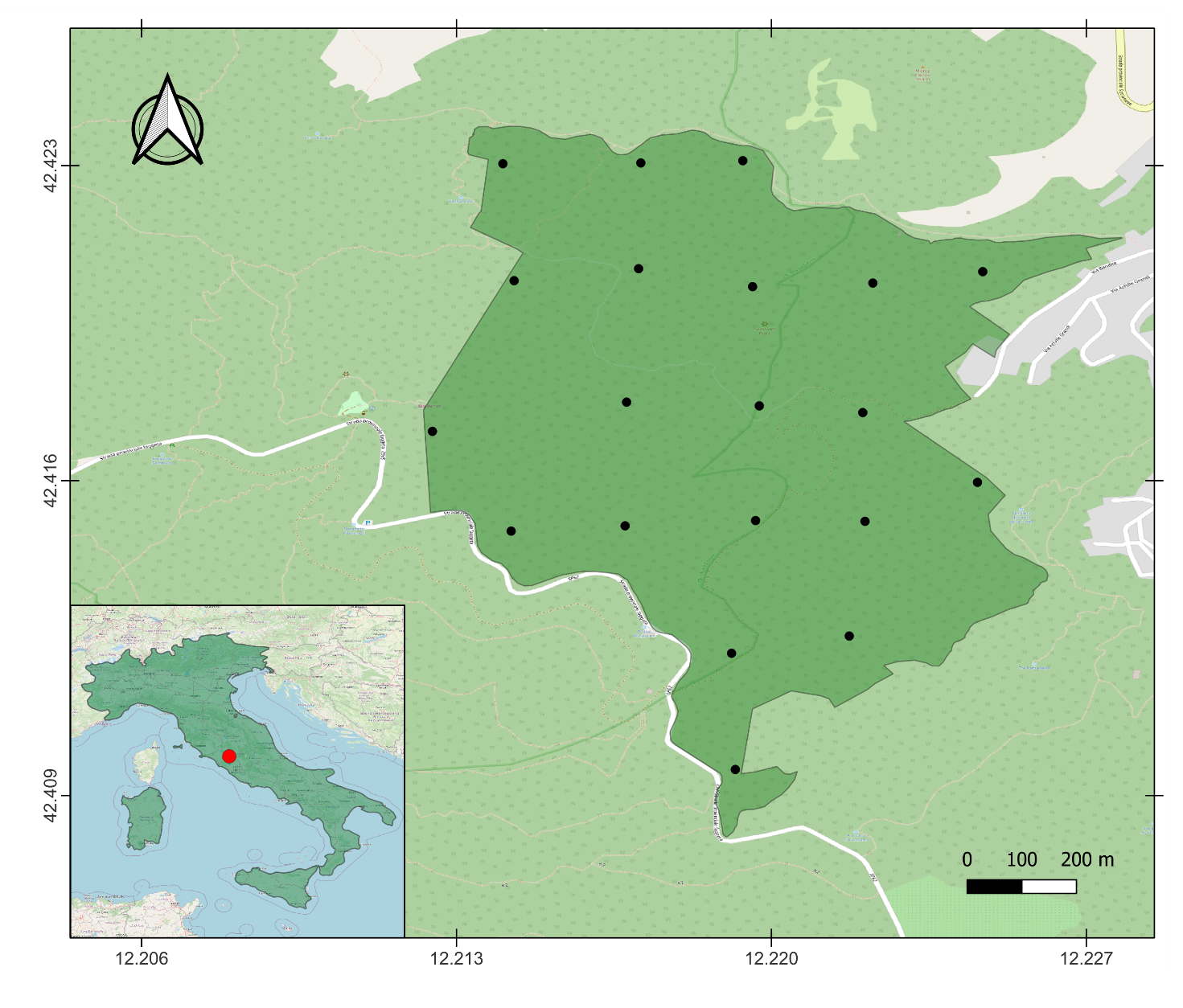
**Figure S1.** Tenuta Sant’Egidio reserve in Mount Cimino, Viterbo, central Italy. The black dots represent the locations of camera-traps within the study area.

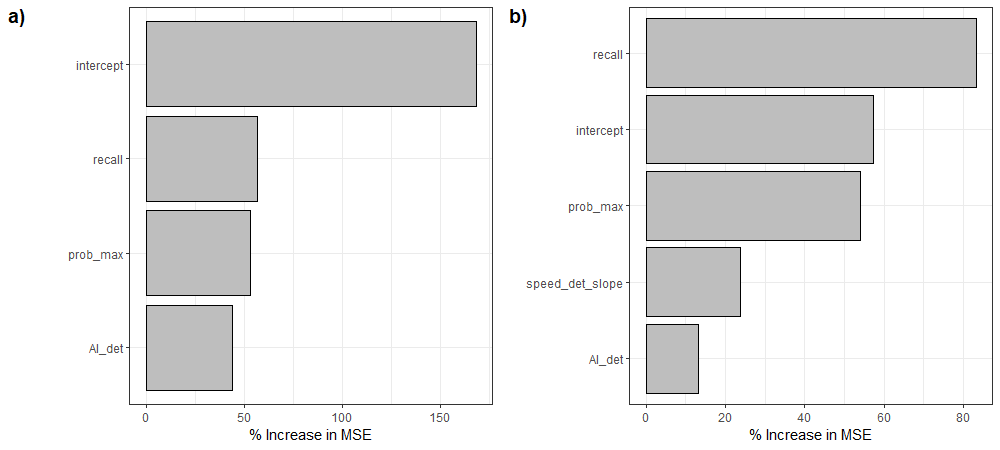

**Figure S2.** Predictive importance of each variable in the random forest models according to the percentage increase in Mean Squared Error (MSE) for a) CT-DS and b) REM. Higher values of %MSE increase are associated with a higher predictive importance of that particular variable in explaining the variation in density when a machine learning algorithm is used to classify images.

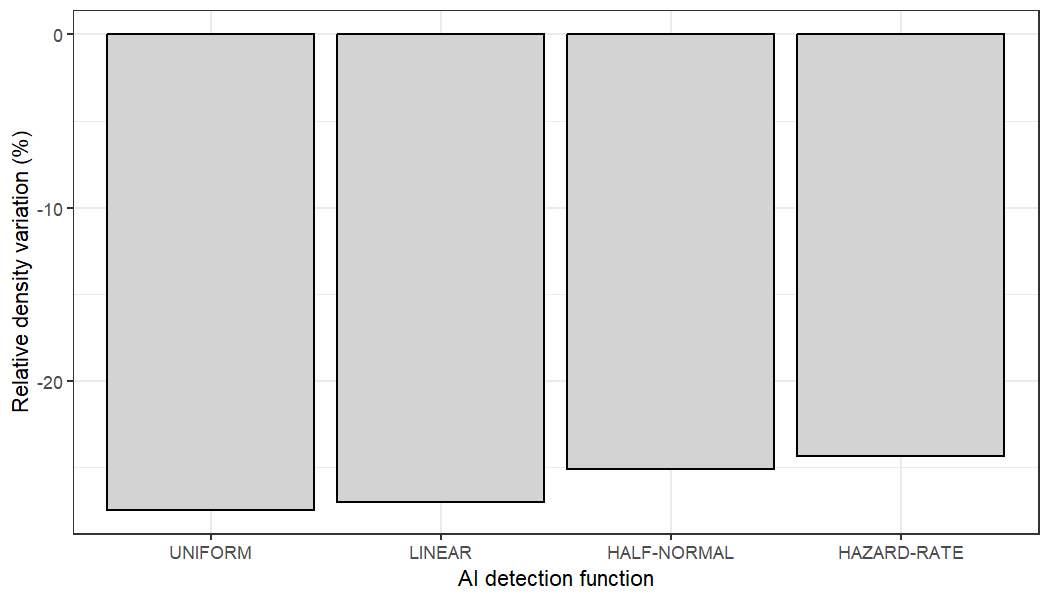

**Figure S3.** Marginal effects on the relative variation in density for the different shapes of the AI detection function considered in the simulations. The half-normal and hazard-rate curves exhibit a slightly better compensation effect, resulting in less underestimation of the density parameter.

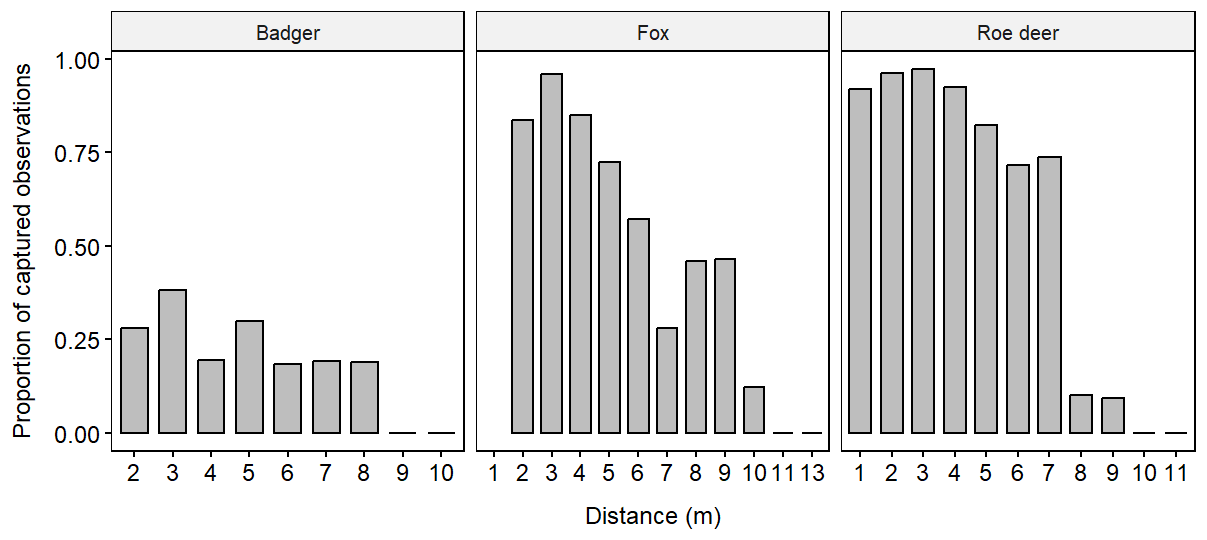
 **Figure S4.** Proportion of observations correctly captured by Wildlife Insights by distance for each focal species. For each class, the proportion is calculated as the number of captured observations over the total number of observations in that class.

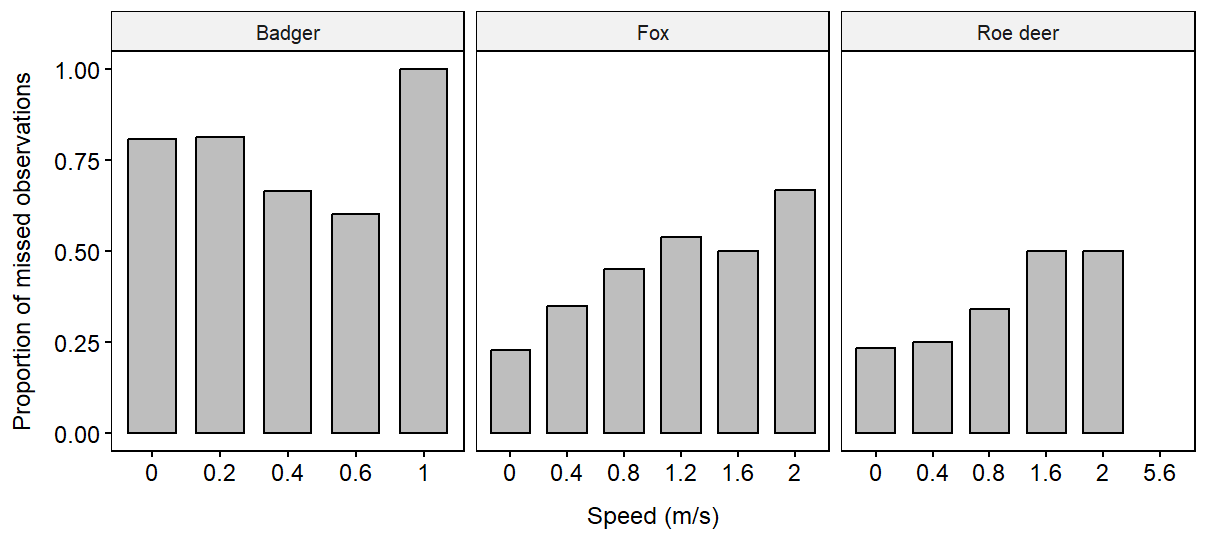

**Figure S5.** Proportion of observations missed by Wildlife Insights by speed class for each focal species. For each class, the proportion is calculated as the number of missed observations over the total number of observations in that class.

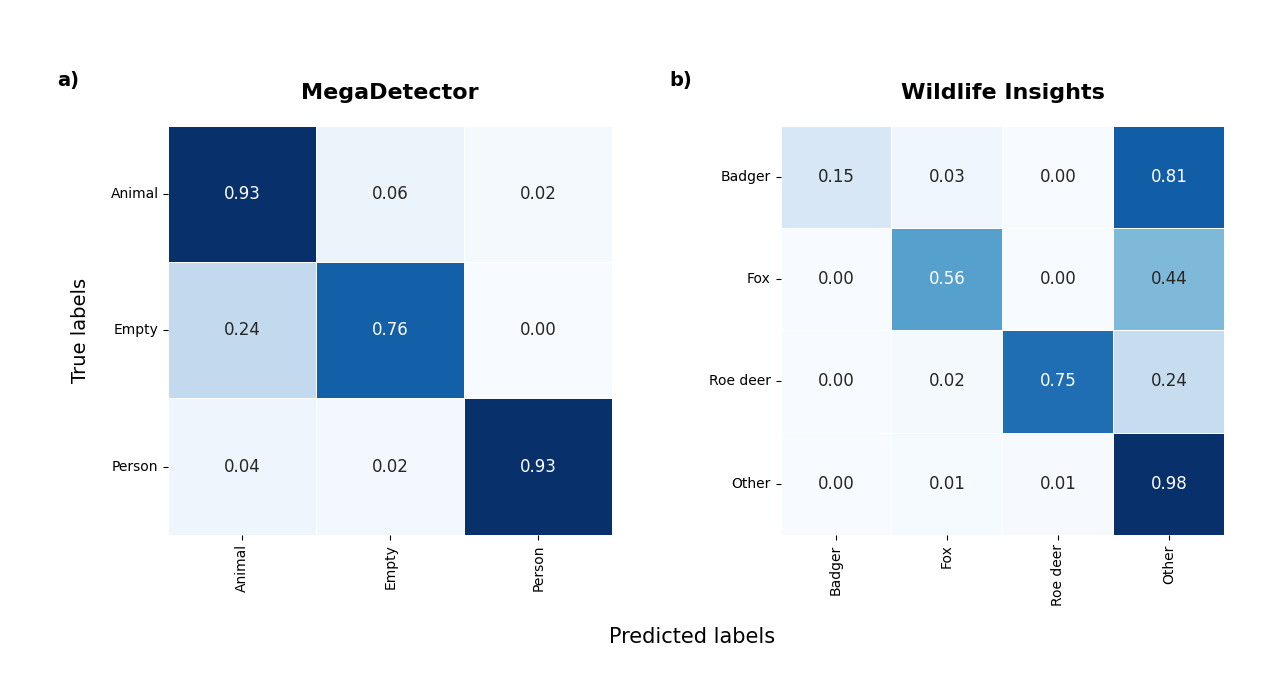

**Figure S6.** Confusion matrices for model performance of a) MegaDetector (animal detection) and b) Wildlife Insights (animal classification). Cells show the proportion of correct classifications for each class against the ground truth, allowing visualization of exclusion errors for that class. “Other” refers to the non-focal animal species captured during the camera-trapping survey.

**Table S1.** Estimated models’ parameters and relative standard errors for REM and CT-DS for the three focal species, both using the user-classified dataset and the WI (Wildlife Insights)-classified dataset. Note that for CT-DS the total capture events differ from the total number of images for that species, due to the fact that t = 1 s was considered for the snapshots so if multiple images were recorded within the same second, only the first was selected.

|  |  | **User image classification** | | | **MD image classification** | | | **WI image classification** | | |
| --- | --- | --- | --- | --- | --- | --- | --- | --- | --- | --- |
| **Method** | **Parameter** | **Badger** | **Red fox** | **Roe deer** | **Badger** | **Red fox** | **Roe deer** | **Badger** | **Red fox** | **Roe deer** |
| CT-DS | Total capture events | 437 | 579 | 523 | 424 | 552 | 514 | 110 | 382 | 394 |
|  | Radius (m) | 6.24 ± 0.14 | 6.01 ± 0.24 | 5.86 ± 0.05 | 6.17 ± 0.15 | 5.93 ± 0.21 | 5.72 ± 0.43 | 5.07 ± 0.30 | 4.35 ± 0.07 | 5.00 ± 0.03 |
| REM | Individuals | 47 | 115 | 67 | 47 | 112 | 67 | 30 | 94 | 57 |
|  | Day range (km/day) | 8.00 ± 1.09 | 9.08 ± 1.48 | 7.82 ± 1.44 | 8.08 ± 1.09 | 10.89 ± 1.12 | 7.13 ± 1.44 | 3.88 ± 0.73 | 6.10 ± 1.30 | 6.36 ± 1.45 |
|  | Radius (m) | 4.56 ± 0.35 | 6.11 ± 0.51 | 5.95 ± 0.39 | 4.57 ± 0.34 | 5.42 ± 0.21 | 5.76 ± 0.40 | 4.06 ± 0.35 | 4.47 ± 0.35 | 3.36 ± 1.20 |
|  | Angle (rad) | 0.64 ± 0.14 | 0.66 ± 0.08 | 0.96 ± 0.00 | 0.64 ± 0.14 | 0.64 ± 0.07 | 0.96 ± 0.00 | 0.64 ± 0.15 | 0.64 ± 0.09 | 0.96 ± 0.00 |
|  | Activity level | 0.40 ± 0.06 | 0.33 ± 0.05 | 0.63 ± 0.10 | 0.40 ± 0.06 | 0.33 ± 0.05 | 0.57 ± 0.10 | 0.30 ± 0.05 | 0.31 ± 0.06 | 0.62 ± 0.10 |
